# Introducing an argonaute-facilitated PCR platform

**DOI:** 10.1101/2022.02.22.481407

**Authors:** Feng Gao, Bojia Han, Yin Chen, Fangfang Sun, Jianhui Yang, Chun-Yu Han

**Affiliations:** Gene Editing Research Center, Hebei University of Science and Technology, Shijiazhuang, Hebei 050018, China

## Abstract

Argonaute proteins are characterized by highly efficient guide DNA/RNA-directed binding to target nucleic acids with high fidelity. Employing this feature, we designed an argonaute-facilitated PCR platform by making use of an argonaute derived from Mesorhizobium japonicum (MejAgo) which does not carry nuclease activity and exposes the 3’ end of guide DNA when binding. Each reaction cycle of the MejAgo-PCR platform consists of a denaturing step and a polymerase-mediated extension step, omitting the annealing step required by the traditional PCR. More importantly, MejAgo-PCR could significantly improve the sensitivity of PCR in template detection due to the argonaute-facilitated pairing between guide DNA/primer and template. Thus, an argonaute-facilitated PCR has the potential to be developed as an advanced PCR platform with higher sensitivity and efficiency.

## Introduction

The invention of polymerase chain reaction (PCR) has significantly advanced modern biology research. In addition, given the fact that the test-trace-and-isolate strategy (TETRIS/TTI) is so far the most efficient way to bring contagious diseases under control, PCR has been widely used in healthcare services. A marked example is the usage of PCR in detecting the infected but asymptomatic individuals in the current COVID-19 pandemic(1). Although many new techniques of nucleic-acid detection have emerged, quantitative PCR (qPCR)-based detection platforms are still considered the gold standard for COVID-19 testing, as recommended by the World Health Organization (W.H.O)(2).

Nucleic acid amplification is the foundation for all the developed nucleic-acid detection systems including the traditional PCR. No exception, the recently developed Specific High Sensitivity Enzymatic Reporter UnLOCKing (SHERLOCK) system makes use of recombinase polymerase amplification (RPA) (3,4). As such, the sensitivity of a certain detection system is determined by the sensitivity of its dependent amplification strategy. In this sense, although the SHERLOCK system is featured with a “collateral effect” of promiscuous RNAse activity, the initial RPA-facilitated amplification determines the reliability of the followed Cas13a-mediated second-order signal amplification. Thus, if the titer of the target template is below the threshold of RPA sensitivity, the claimed advantage of SHERLOCK is rootless.

The traditional PCR is a heterothermal reaction using repeated cycles of heating and cooling to make many copies of a specific region of DNA. In contrast, the isothermal amplification occurs in a stable temperature, which, obviously, has many advantages by being equipment-free, time-saving and portable. The RPA-facilitated PCR is a type of isothermal reaction in which the UvsX recombinase carries the primers and facilitates its pairing with double-stranded DNA (dsDNA) templates. RPA employs recombinase-primer complexes to scan double-stranded DNA and facilitate strand exchange at cognate sites, sparing the high-temperature denaturation process otherwise required by the traditional PCR, so that isothermal amplification could proceed. However, the RPA platform additionally requires single-stranded DNA binding protein which interacts and prevents the displaced template strand from the ejection of the primer by branch migration. Moreover, the recombinase reaction requires the presence of ATP for energy input.(5) All of these supplements talked above increase both the complexity and the expense of application. Moreover, recombination is a complex process with mechanisms not fully addressed(6). As such, no definite rules has been established for optimization of primers used for the RPA platform. Therefore, an additional process of primer screening is unavoidable when RPA is to be applied, which is obviously laborious. Besides, there is no implication that RPA could improve sensitivity of nucleic acid detection as an outcome(5).

No matter is heterothermal or isothermal the amplification process, binding between primers and templates is the first step of amplification and the efficacy of binding determines the sensitivity of the followed amplification. The primer-template pairing process of the traditional PCR is totally dependent on random physical interaction between primers and templates. In such a thermodynamic laws-relied system, increasing the concentration of primers could definitely increment the efficiency of pairing with templates. However, due to the non-specific mispairing between primers and between primers and templates, a trade-off effect on accuracy has to be considered.

Argonaute proteins are characterized by highly efficient guide DNA/RNA-directed binding to target nucleic acids with high fidelity. Researches on mechanisms of argonaute-facilitated targeting revealed that argonaute proteins could accelerate guide binding to target nucleic acids(7,8,9,10). In this study, we demonstrated that an argonaute derived from Mesorhizobium japonicum (MejAgo) could be employed to enhance primer-template pairing. With this feature at play, a MejAgo-facilitated PCR system could improve sensitivity in nucleic acid detection.

## Results

An ideal argonaute suitable for application in PCR should carry two features. First, the 3’ end of single-stranded guide DNA (gDNA) must be free of occupation by the argonaute protein (Fig. 1) since DNA polymerase acts on the 3’-OH of the existing strand (primer in this case) for adding free nucleotides. We predict that a nonclassical argonaute lacking the PAZ domain would satisfy this purpose since the PAZ domain is responsible for argonaute binding with the 3’ end of gDNA(11). Second, the argonaute must be inactive in nuclease activity. It is known that several highly conserved amino acid residues, i.e., DEDX (X stands for D or H) (11), are required for the nuclease activity of classical argonautes. Thus, an argonaute with mutations at these residues is preferred. By screening of prokaryotic argonautes (most eukaryotic argonautes bind to RNA instead), we isolated an argonaute derived from Mesorhizobium japonicum (MAFF 303099; GenBank: BAB52533.1) (designated as MejAgo) which satisfies both requirements (Fig. 2).

**Figure 1.**
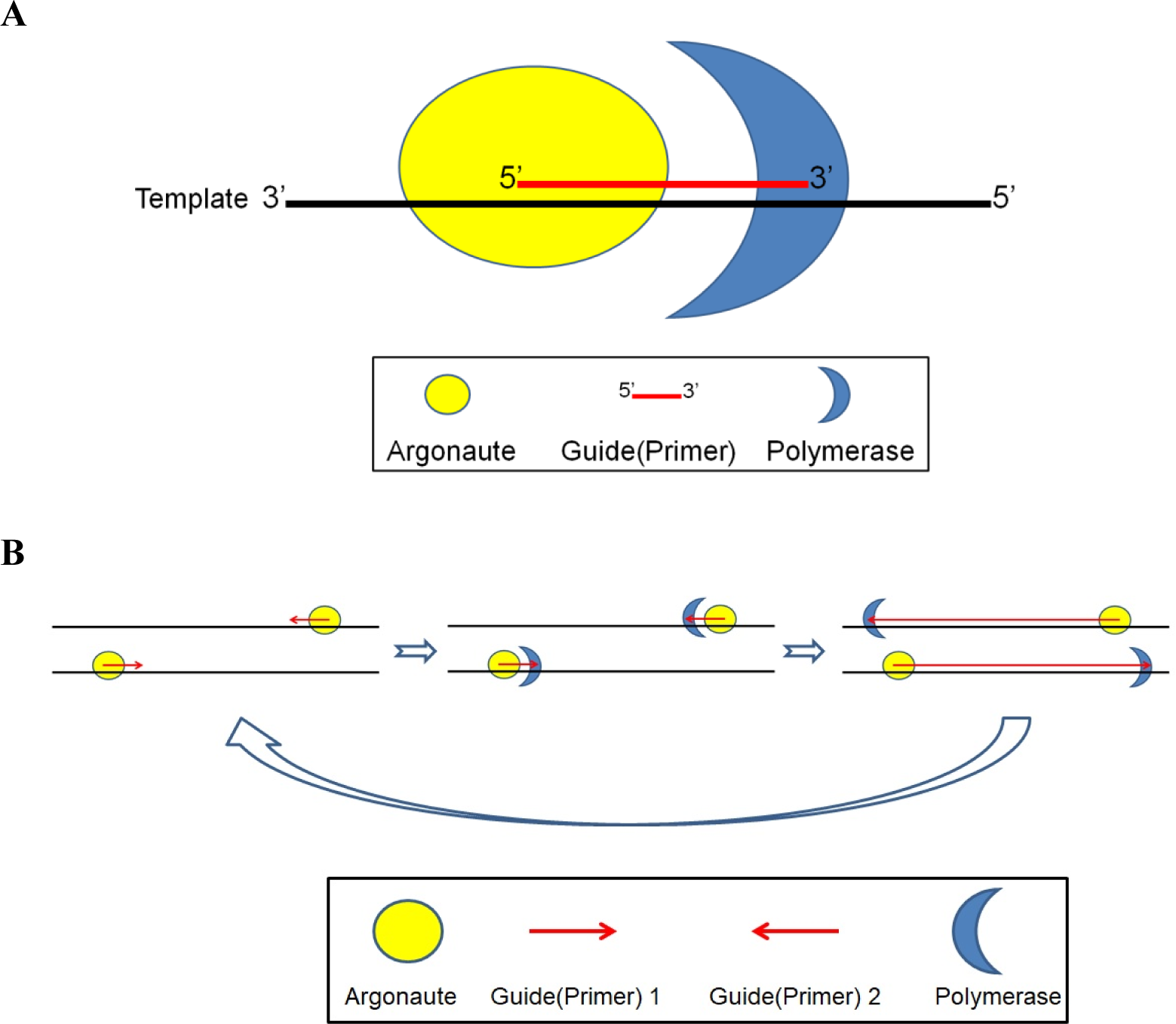
**(A)** A strategic model for an argonaute-involved primed polymerization, in which the 3’ end of the guide/primer should be exposed for reaction with DNA polymerases. **(B)** The theoretical working model for an argonaute-facilitated PCR platform.

**Figure 2.**
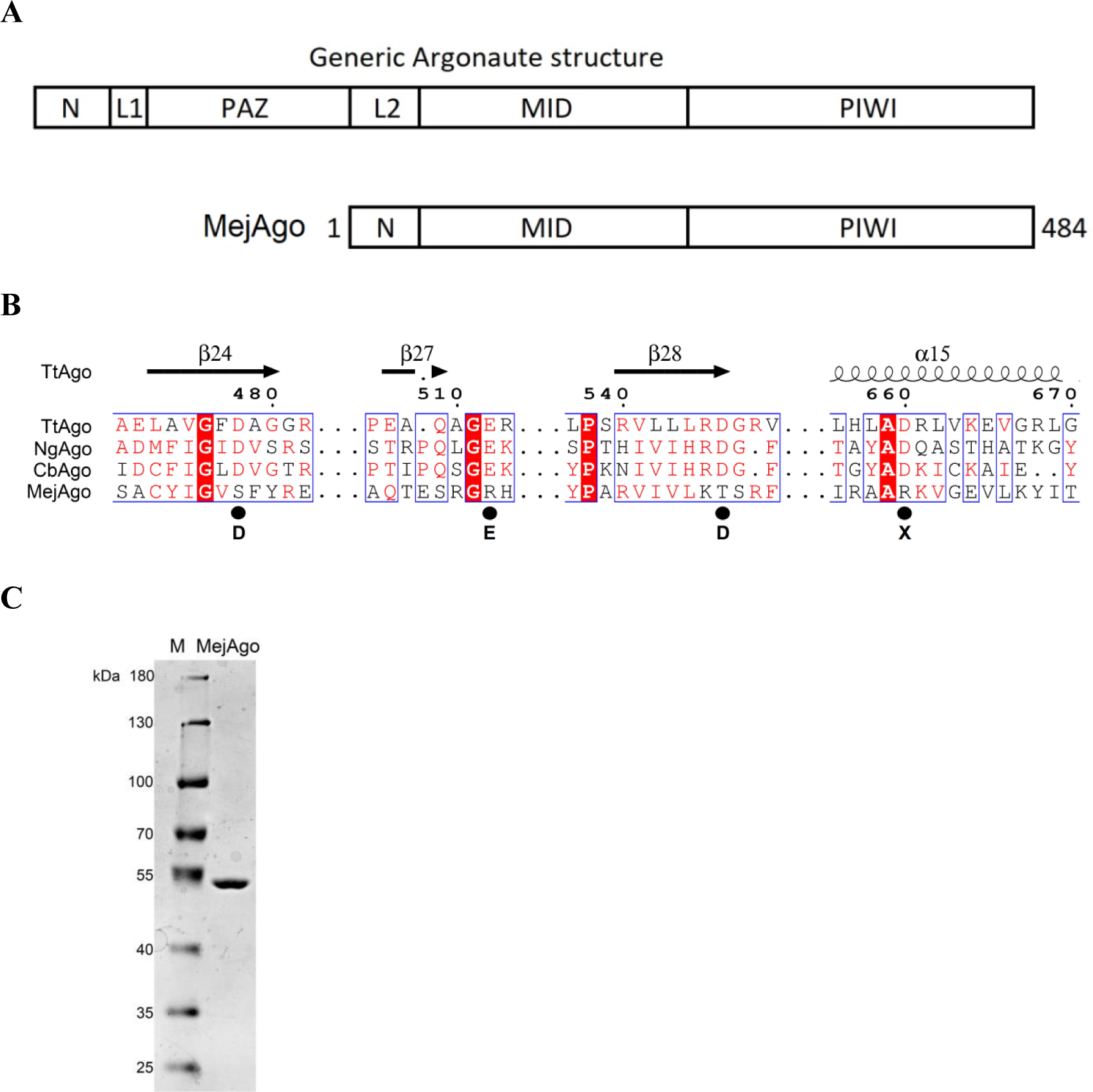
The structural characteristics of MejAgo. **(A)** Schematics of domain organization of a generic argonaute protein and the MejAgo. **(B)** Sequence alignment of the PIWI domains of MejAgo, TtAgo, NgAgo, and CbAgo. The MejAgo has mutations at the residues (DEDX) critical for nuclease activity. **(C)** PAGE analysis of the purified recombinant MejAgo.

A traditional PCR procedure consists of a high temperature (above 90 °C) denaturation process, an annealing process with relatively low temperature (below 60 °C) followed by a Taq polymerase-mediated extension reaction (∼70 °C). To demonstrate the proof of concept that argonaute could facilitate primer-template pairing (annealing), we modified the conditions and the procedure of traditional PCR based on the characteristics of MejAgo: 82 °C was selected as the denaturing temperature since MejAgo could not tolerate a temperature above 90°C, followed by Taq polymerase-mediated extension at 68°C, omitting the annealing process (Fig. 3A). To amplify a sequence of M13 lac phage vector M13mp18, we chose a pair of gDNAs (functioning as primers), one complementary to the 5’ end of this sequence and the other aligned to the 3’ end of this sequence. Indeed, with the modified protocol, addition of MejAgo could remarkably enhance the amplification of the target DNA (Fig. 3B-D).

**Figure 3.**
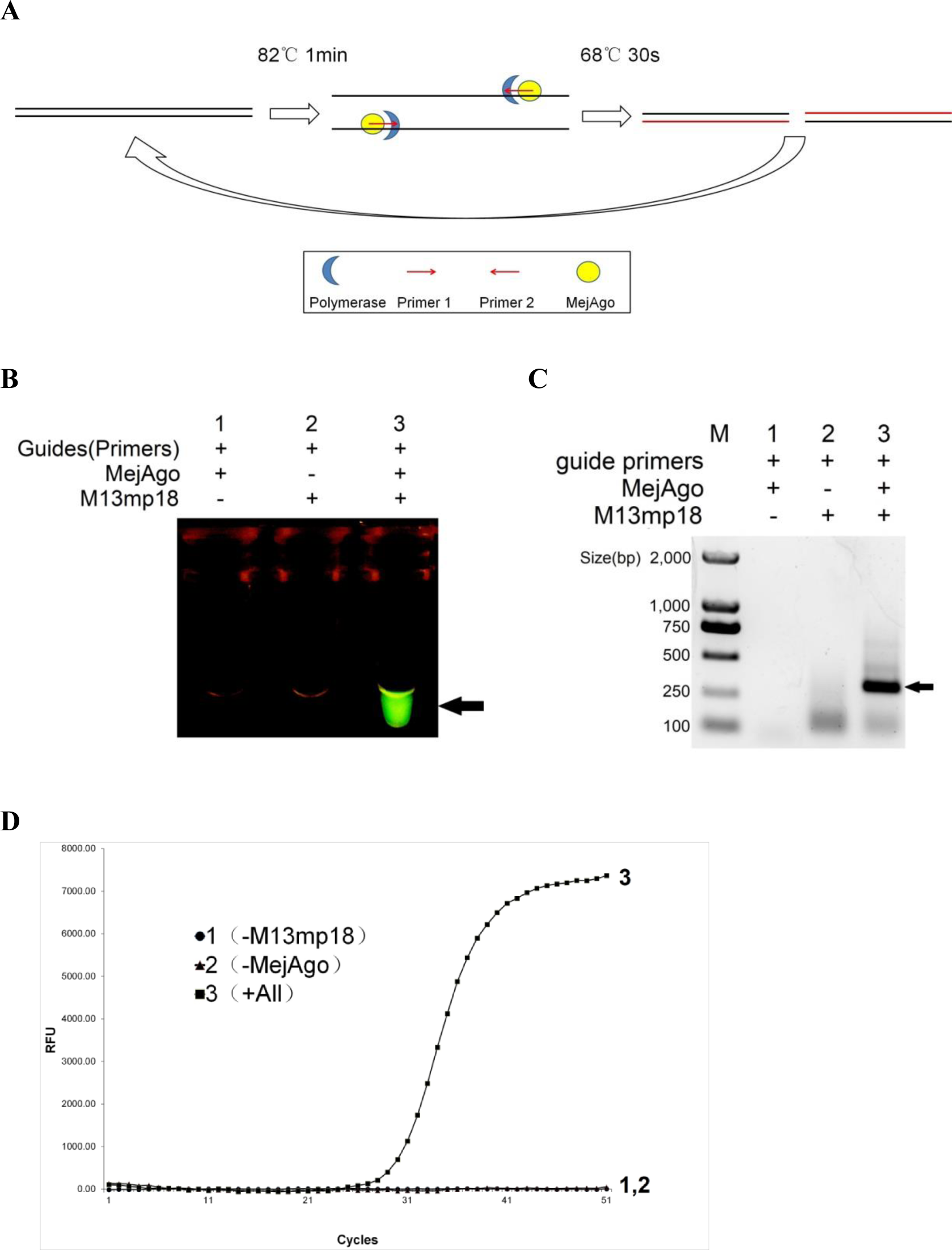
Demonstration of the MejAgo-PCR platform. **(A)** A working diagram of the two-step program of the MejAgo-PCR platform. **(B-D)** Amplification of the target sequence on the M13mp18 vector was examined by (B) 470nm blue light, (C) argrose electrophoresis, and (D) real-time fluorescence detection. 6×10^5^ copies/20µl of M13mp18 template was used in these experiments. The guides/primers used in these experiments are pM13-3-50 and pM13-25-50. The probe used for qPCR is F-M13-2. Sequences of primers and the probe are detailed in Materials and Methods.

Not only for the fragments of single-stranded circular DNA, but sequences of linear double-stranded DNA and double-stranded circular plasmid DNA could also be successfully amplified by this MejAgo-PCR platform, verifying its broad application (Fig. 4).

**Figure 4.**
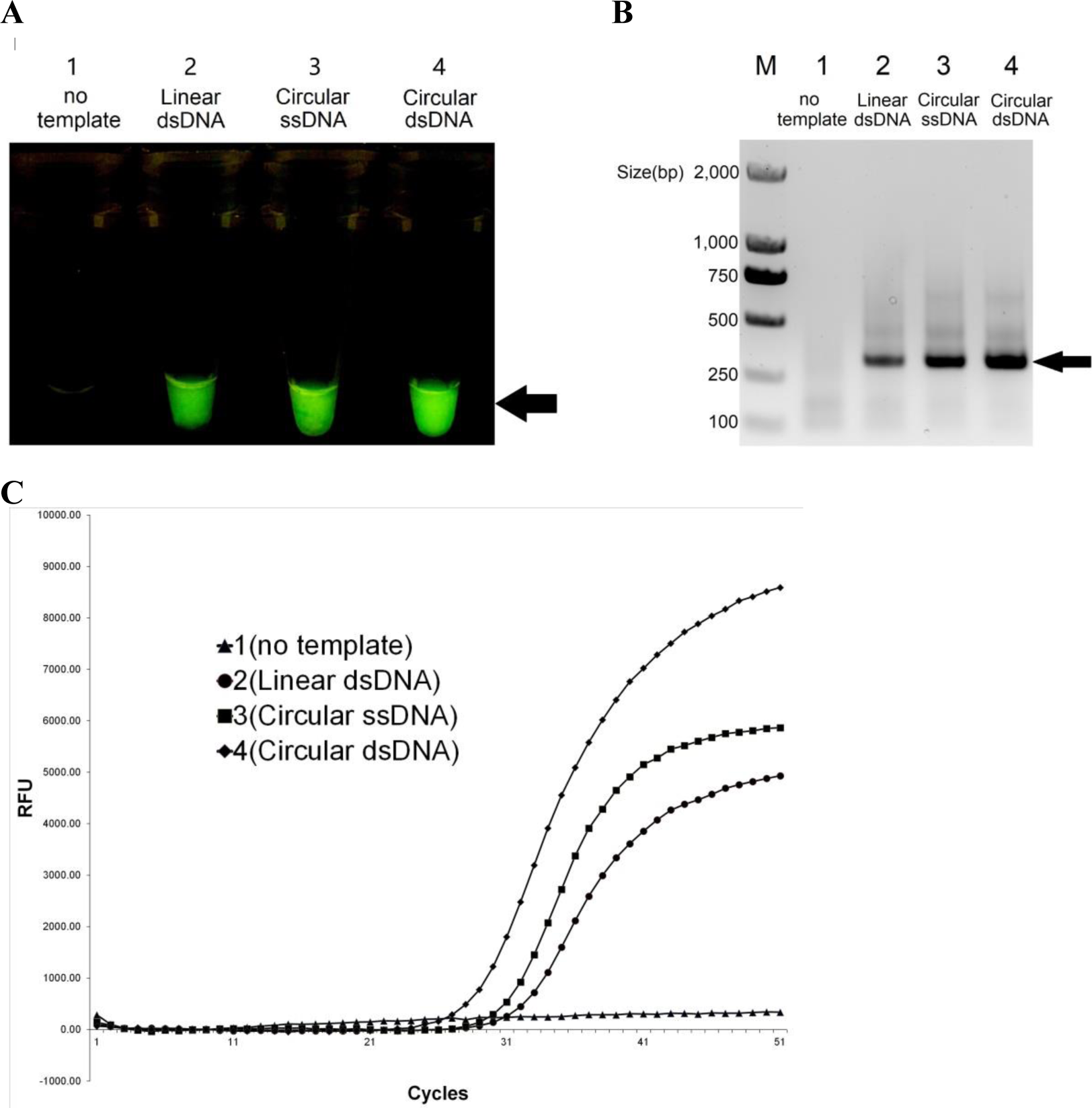
MejAgo-PCR is capable of amplifying various DNA substrates. M13mp18 vector-based linear dsDNA(dsDNA1300), circular single-stranded DNA(M13mp18) and circular double-stranded DNA(pM13 plasmid) templates were tested. Resultant products were analyzed by **(A)** 470nm blue light, **(B)** argrose electrophoresis, and **(C)** real-time fluorescence detection. The guides/primers used in these experiments are pM13-3-50 and pM13-25-50. The probe used for qPCR is F-M13-2. Sequences of primers and the probe are detailed in Materials and Methods.

Length of gDNAs is a critical variable for PCR. We tested gDNAs ranging 10-50 nt and found that ⩾27 nt is preferable by the MejAgo-PCR (Fig 5), as shorter guides sharply reduced the efficiency (data not shown), possibly due to the free-3’-end requirement talked above.

**Figure 5.**
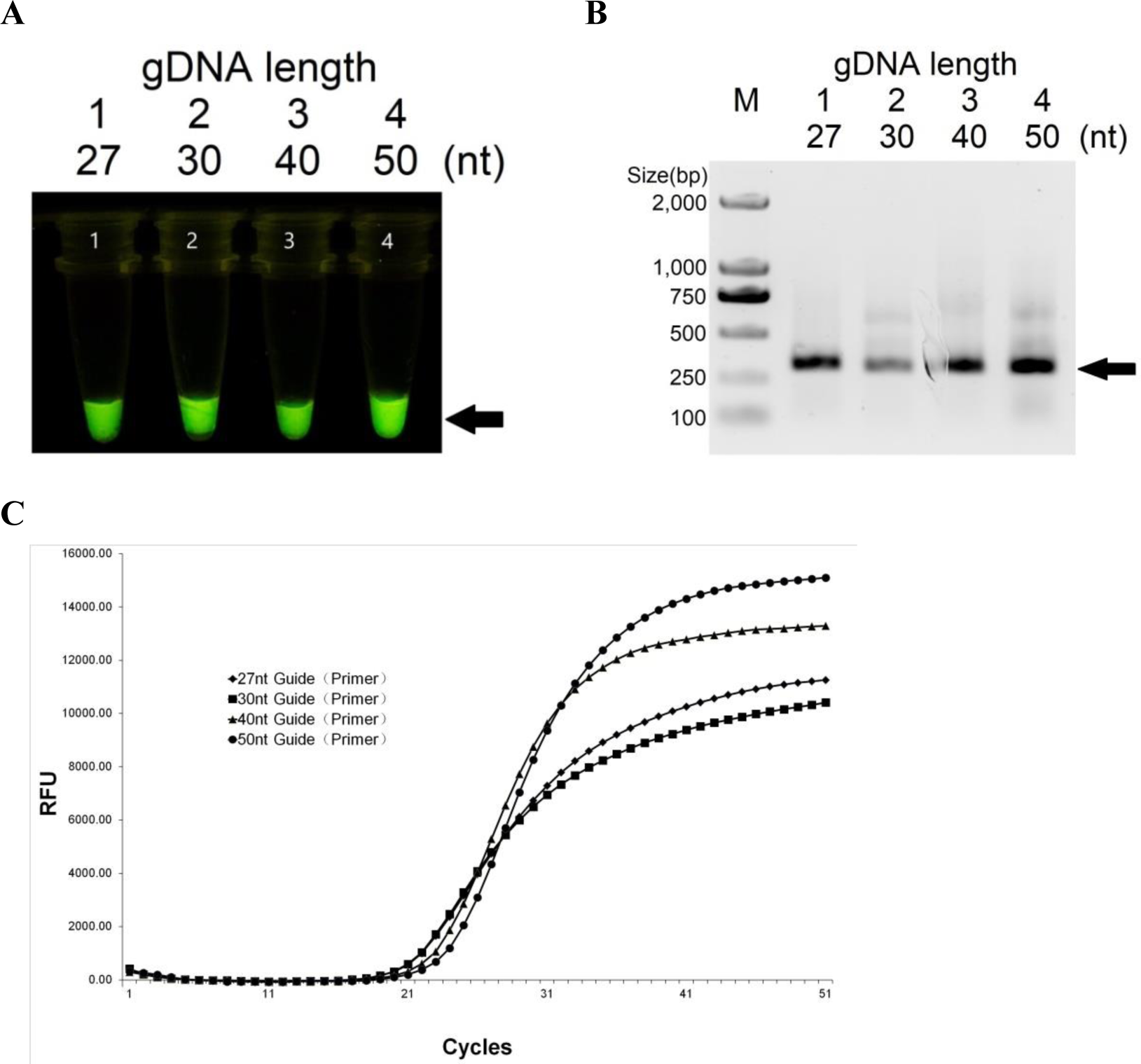
Primers with ≥27nt length is preferable for the MejAgo-PCR platform. Different lengths of primers targeting the same sequence of the M13mp18 vector as Figures 3 and 4. Resultant products were analyzed by **(A)** 470nm blue light, **(B)** argrose electrophoresis, and **(C)** real-time fluorescence detection. 6×10^6^ copies/20µl of M13mp18 template was used in these experiments. Primers used in the experiments are pM13-3-27, pM13-3-30, pM13-3-40, pM13-3-50 and pM13-25-27, pM13-25-27, pM13-25-27, pM13-25-50. The probe for qPCR is F-M13-2. Sequences of primers and probe are detailed in Materials and Methods.

Omitting the annealing process from the traditional PCR is beneficial from shortening the time required for a PCR cycle. Another posited advantage of the MejAgo-PCR over traditional PCR is based on the fact that argonautes could accelerate gDNA (primer) binding to target (template) (7,8,9,10). Therefore, we hypothesized that the MejAgo-PCR platform could be more sensitive than the traditional one in detecting template signals. To probe this possibility, we ran a series of PCR tests for a sequence of the M13mp18 vector with serial dilutions of template copies. As shown in Fig. 6, within the same reaction cycles, the MejAgo-PCR platform was able to detect as few as 6 copies of the template, two orders of magnitude improvement in sensitivity over the traditional PCR platform.

**Figure 6.**
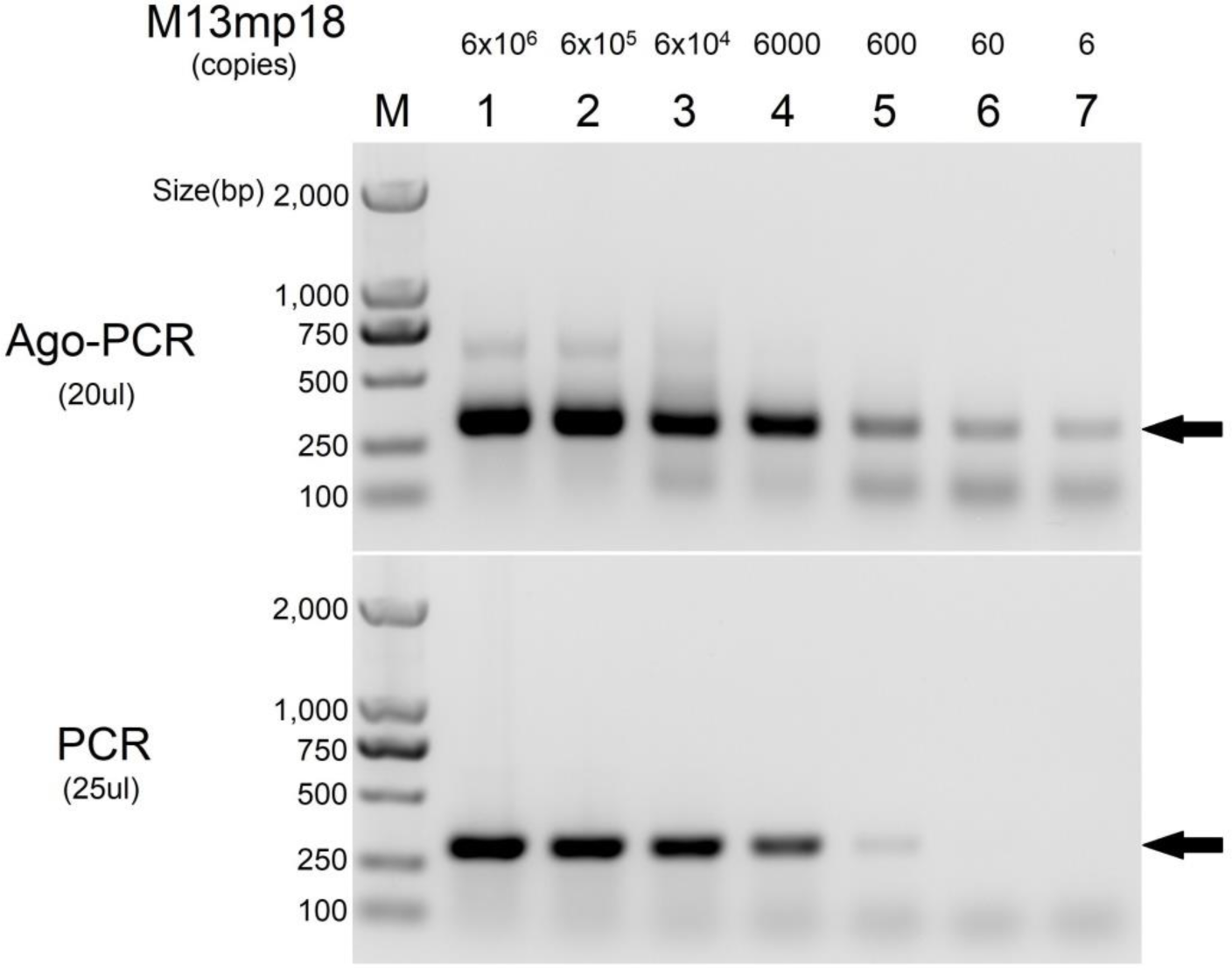
Comparison of the MejAgo-PCR platform with the traditional PCR platforms in sensitivity. The PCR reaction was a standard 3-step program (95°C 3min, 95°C 30s, 57°C 30s, 68°C 30s, step 5 Go to step 2,40 cycles, 68°C 5min incubation, end) with a pair of optimized primers (oM13-3-40 and oM13-25-30). The MejAgo-PCR reaction was a 2-step program (82°C 1min, 68°C 30s, 40cycles, end). The primers used in the MejAgo-PCR reaction were pM13-3-50 and pM13-25-50. Sequences of primers are detailed in Materials and Methods. The ssDNA template M13mp18 was serially diluted to the final concentrations as indicated on top of each line.

At last, we examined the application of the MejAgo-PCR platform to detect target sequence in an environment of genomic DNAs, a condition when PCR was more often applied. As assessed by the linearity of the quantification cycle (Cq) curve, the MejAgo-facilitated PCR platform could again detect 6 copies of M13mp18 template admixed in 300 ng mammalian genomic DNAs (Fig. 7). In conclusion, the above results manifest that MejAgo-PCR has the potential to be a more efficient DNA detection platform.

**Figure 7.**
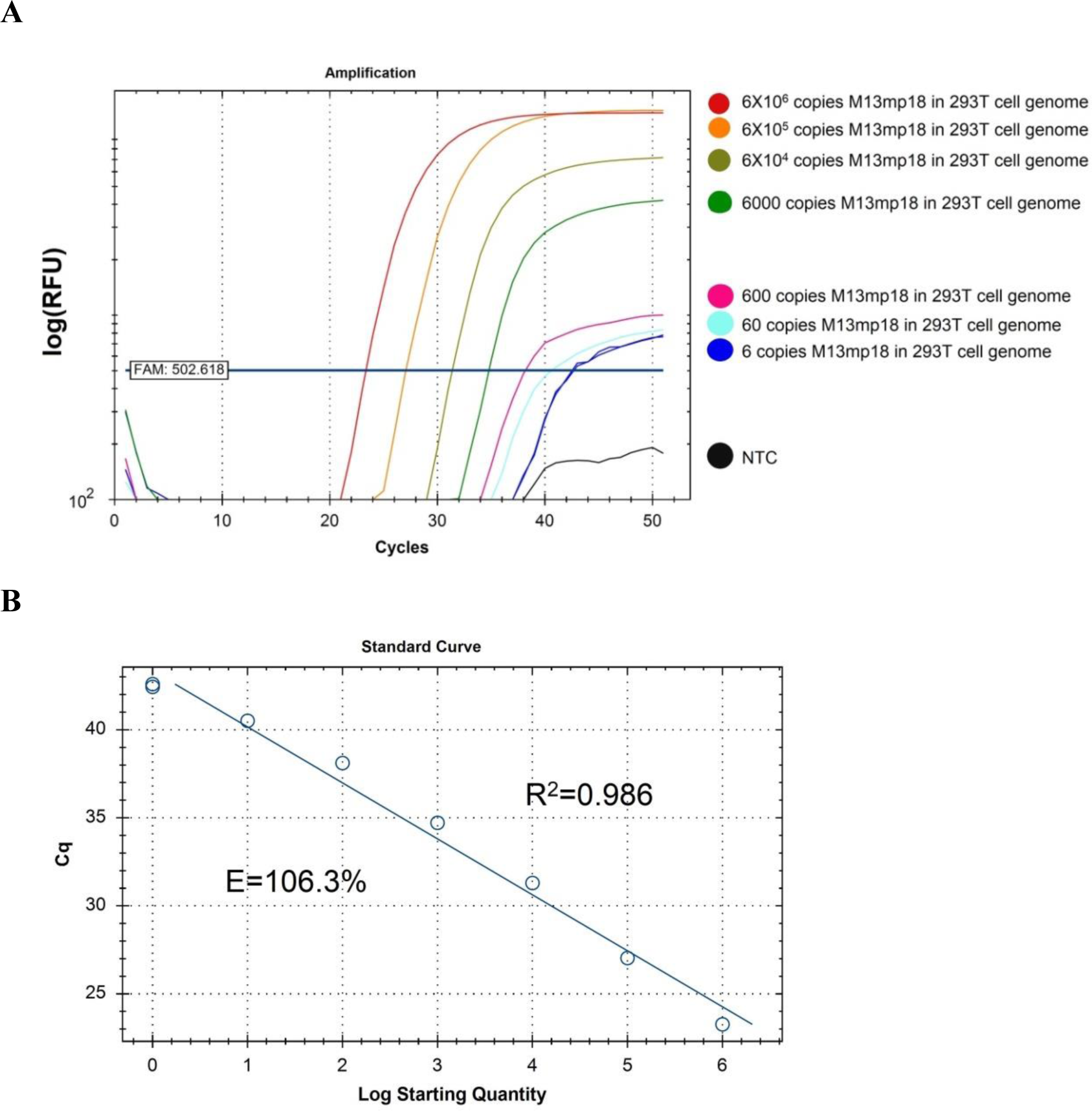
MejAgo-PCR is superior to the traditional PCR in detecting low-copy target in genome DNA-mixed samples. The target M13mp18 vector (template) was serially diluted, as indicated, to the samples with 300 ng genomic DNA extracted from 293T cells. Final concentrations of template are shown as copies per tube (20 µl). **(A)** The resulting products is shown by the real-time fluorescence detection curve. **(B)** The cq curve analysis validates the successful detection of template in as low as 6 copies. The primers used in the experiments were pM13-3-50 and pM13-25-50. The probe for qPCR was F-M13-2. Sequences of guides and probe are detailed in Materials and Methods.

## Discussion

In this study, we demonstrated that the MejAgo could be utilized as a tool to enhance the traditional PCR platform in the sensitivity of nucleic acid detection, proving our hypothesis that an argonaute protein which lacks both the PAZ domain and nuclease activity could be applicable in PCR for facilitating primer-template pairing. Argonautes widely exist in the biological world. Besides the DNA-targeting argonautes, other argonautes from some species preferably target RNAs. It is thus conceived that some argonautes could be developed for reverse-transcriptase PCR (RT-PCR). In addition, it is feasible that with the assistance of appropriate DNA helicase, argonaute-based isothermal PCR system could be developed.

With the assistance of argonaute in primer-template pairing, the annealing step of the traditional PCR reaction could be omitted in the Ago-PCR platform, thus saving the time of amplification. Also, the MejAgo-facilitated pairing is known to function in a wide range of temperatures, suggesting its suitability in conjunction with other DNA polymerases besides Taq polymerase. More importantly, we demonstrated that the Ago-PCR platform was more sensitive than the traditional platform, consistent with the previous reports showing that argonaute could increase the efficiency of pairing of guide DNAs/RNAs with nucleic acid targets (7,8,9,10).

## ACKNOWLEDGEMENTS

We thank professor Xiao Z. Shen (Department of Physiology, Zhejiang University School of Medicine, Hangzhou, Zhejiang, China.) for rewritting the manuscript.

## Materials and Methods

### Protein expression and purification

For expressing MejAgo(Mesorhizobium japonicum MAFF 303099,GenBank: BAB52533.1).MejAgo CDS was cloned into pET-28a(+) and transformed in BL21(DE3). Recombinant MejAgo protein expression and Ni-affinity chromatography purification procedures were performed according to user manual(Novagen). In perticular, the resultant purified MejAgo protein was eluted with elution buffer(50 mmol/L Tris-HCl pH 7.5, 500 mmol/L NaCl, 1 mmol/L DTT, 250 mmol/L Imidazole), and concentrated with 50Kda colum(Merck Millipore#UFC905008) to 1µg/µl. Adding glycerol to final concerntration of 50% for preservation.

### MejAgo-PCR

For MejAgo PCR, 900nM MejAgo protein, 200nM foward guide/primer(pM13-3-50) and 200nM reverse guide/primer(pM13-25-50) were preincubated at 55°C in Ago-PCR buffer(20 mM Tris-HCl, 10 mM (NH4)2SO4, 50 mM KCl, 4 mM MgSO4, 0.1% Tween®20, 100ug/ml BSA, pH 8.8) for 15 min, then add DNA template, 250uM dNTP Mixture, 1U Taq DNA Polymerase(NEB), add water to final volum to 20µl. MejAgo-PCR program: 82°C 1min,68°C 30s,40 cycles. 1% argose electrophoresis analysis for Ago-PCR products.

For MejAgo qPCR, reaction buffer and system are the same to Ago-PCR, except that 250nM Taqman probe F-M13-2 was added along with Taq polymerase, dNTP, DNA template. Seal the reaction mixture with 10µl mineral oil. qPCR(BIO-RAD CFX-96) program: 82°C 1min,68°C 30s,snap;50 cycles.

### PCR

For tranditional PCR, Primers against the same region of MejAgo-PCR is optimized with length according to the commonly used annealing temperature of a PCR program. Reaction system: 200nM foward primer(oM13-3-40), 200nM reverse primer(oM13-25-30), 200µM dNTP, 1U Taq DNA Polymerase(NEB), 1X Standard Taq Reaction Buffer(NEB B9014S) and certain dosage of M13mp18 template according needs. Program: 95°C 3min, 95°C 30s, 57°C 30s, 68°C 30s, step 5 Go to step 2, 40 cycles, 68°C 5min end.

### Template prepration

The 4330bp PCR product amplified from M13mp18 by using primers: oM13-5-27 and oM13-3-24 was inserted into pGEM®-T Easy(PROMEGA #A137A). The resultant plasmid is named as pM13. pM13 is used as double stranded, circular template. The double stranded, linear template(dsDNA1300) was produced by amplifying M13mp18 by using primers oYC-299 and oYC-300. The resultant PCR product is 1300bp.

### Sequences of guides and primers

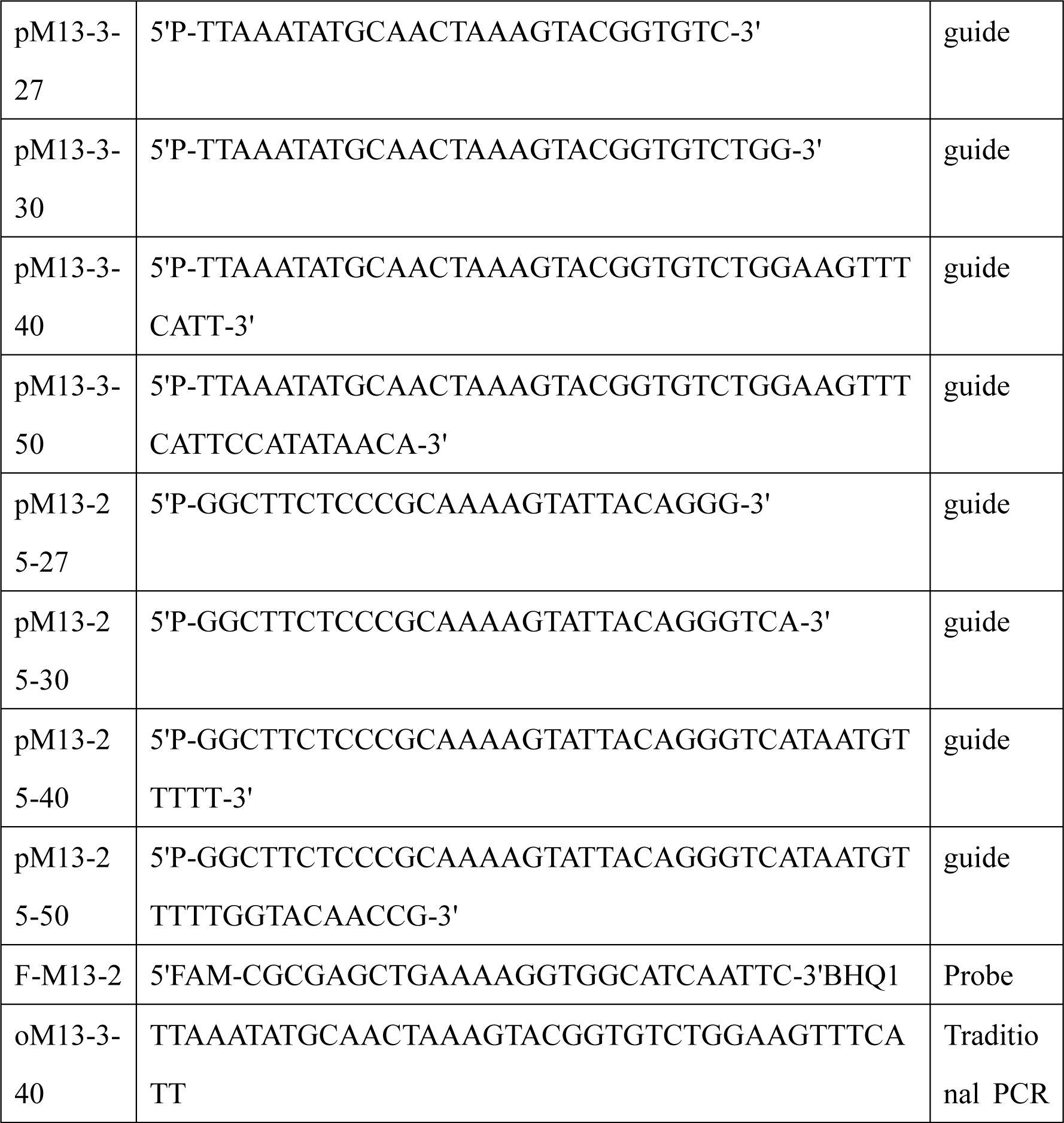
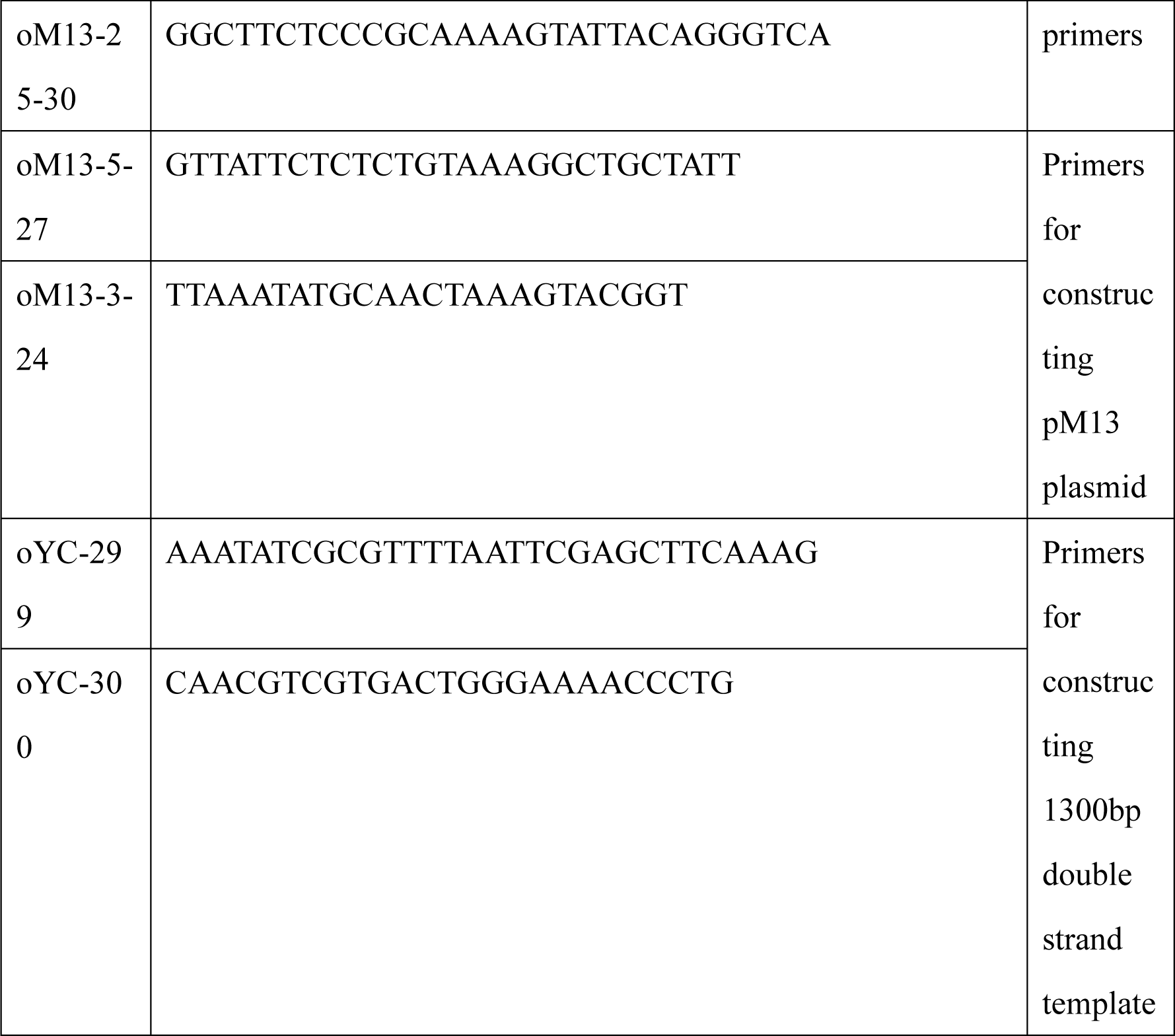

